# Locus cœruleus noradrenergic neurons phase-lock to prefrontal and hippocampal infra-slow rhythms that synchronize to behavioral events

**DOI:** 10.1101/2022.05.12.491630

**Authors:** Liyang Xiang, Antoine Harel, Ralitsa Todorova, HongYing Gao, Susan J. Sara, Sidney I. Wiener

**Affiliations:** Center for Interdisciplinary Research in Biology (CIRB), College de France, CNRS, INSERM, PSL Research University, Paris, France; Zhejiang Key Laboratory of Neuroelectronics and Brain Computer Interface Technology, Hangzhou, China; Department of Child and Adolescent Psychiatry, New York University Medical School, New York, NY, USA

**Author notes:** **Correspondence:** Sidney Wiener.

**Keywords:** phase reset, local field potentials, oscillations, integrative function, chronic recordings

## Abstract

The locus cœruleus (LC) is the primary source of noradrenergic projections to the forebrain, and, in prefrontal cortex, is implicated in decision-making and executive function. LC neurons phase-lock to cortical infra-slow wave oscillations during sleep. Such infra-slow rhythms are rarely reported in awake states, despite their interest, since they correspond to the time scale of behavior. Thus, we investigated LC neuronal synchrony with infra-slow rhythms in awake rats performing an attentional set-shifting task. Local field potential (LFP) oscillation cycles in prefrontal cortex and hippocampus on the order of 0.4 Hz phase-locked to task events at crucial maze locations. Indeed, successive cycles of the infra-slow rhythms showed different wavelengths, as if they are periodic oscillations that can reset phase relative to salient events.

Simultaneously recorded infra-slow rhythms in prefrontal cortex and hippocampus could show different cycle durations as well suggesting independent control. Most LC neurons (including optogenetically identified noradrenergic neurons) recorded here were phase-locked to these infra-slow rhythms, as were hippocampal and prefrontal units recorded on the LFP probes. The infra-slow oscillations also phase-modulated gamma amplitude, linking these rhythms at the time scale of behavior to those coordinating neuronal synchrony. This would provide a potential mechanism where noradrenaline, released by LC neurons in concert with the infra-slow rhythm, would facilitate synchronization or reset of these brain networks, underlying behavioral adaptation.

## 1 Introduction

The brain coordinates activity among interconnected regions via coherent oscillatory cycles of excitation and inhibition (Womelsdorf, et al., 2007). This can facilitate communication among selected subsets of neurons, groups of neurons, and brain regions. Sensory stimuli or behavioral events can reset the phase of these oscillations (Canovier, 2016; Voloh and Womelsdorf, 2016), linking activity of multiple neurons to process information in concert. However, the principal brain rhythms studied in behaving animals are at the time scale of cell neurophysiological processes, which are much faster (on the order of tens and hundreds of milliseconds) than real life behavioral events, which typically occur at second and supra-second time scales. The brain has several mechanisms linking these two time scales, some of which involve the hippocampus (reviewed in Banquet, et al., 2021) and associated networks, including the prefrontal cortex and striatum.

Little is known about brain rhythms that operate in this crucial behavioral time scale during awake behavior. The brain is indeed capable of generating rhythms on the order of 0.1-1.0 Hz, although these have been principally characterized during sleep (Steriade, 1993). Furthermore, during sleep or under anesthesia, rat noradrenergic locus cœruleus (LC) and prefrontal cortical (Pfc) neurons are phase-locked to infra-slow rhythms (Lestienne, et al., 1997; Eschenko, et al., 2012; Totah, et al., 2018). LC stimulation exerts powerful influence on neurophysiological activity in Pfc and hippocampus (Hip; Berridge and Foote, 1991). LC actions in prefrontal cortex are implicated in vigilance, decision-making, and executive function, while in Hip they are associated with learning and processing contextual information (e.g., Wagatsuma, et al., 2018; Sara, 2009 for review). Since oscillations can coordinate activity in brain networks, we reasoned that there might also be rhythmicity on this behavioral time scale in awake animals, and investigated this possibility in rats performing a task engaging Pfc, Hip and LC (Oberto, et al., 2022; Xiang, et al., 2019). Such coordinated activity could provide a possible link between neuromodulation and oscillatory coordination of brain areas on the time scale of behavior.

## 2 Materials and Methods

All experiments were carried out in accordance with local (Comité d’éthique en matière d’expérimentation animale no. 59), institutional (Scientific Committee of the animal facilities of the Collège de France) and international (US National Institutes of Health guidelines; Declaration of Helsinki) standards, legal regulations (Certificat no. B751756), and European/national requirements (European Directive 2010/63/EU; French Ministère de l’Enseignement Supérieur et de la Recherche 2016061613071167) regarding the use and care of animals. The data analyzed here were recorded in experiments described by Xiang, et al. (2019) and further details can be found there.

### 2.1 Animals

Nine male Long-Evans rats (Janvier Labs, Le Genest-Saint Isle France; weight, 280–400 g) were maintained on a 12 h:12 h light-dark cycle (lights on at 7 A.M.). The rats were handled on each workday. To motivate animals for behavioral training on the T maze, water was partially restricted except for a 10–30 min period daily to maintain body weight at 85% of normal values according to age. Rats were rehydrated during weekends. Food was restricted to 14 g of rat chow daily (the normal daily requirement) to prevent the animals from becoming overweight. Recording properties and behavioral correlates of the LC neurons of four of the rats were reported in Xiang, et al. (2019), but no LFP data are presented there. Recordings of LC neurons during task performance was not possible in the remaining five rats included here.

### 2.2 The automated T maze with return arms

The behavioral task took place in an elevated automated T-maze (see Fig. 3A) consisting of a start area, a central arm, two reward arms and two return arms, which connected the reward arms to the start area. Small wells at the end of each reward arm delivered liquid reward (30 µl of 0.25% saccharin solution in water) via solenoid valves controlled by a CED Power1401 system (Cambridge Electronic Design, Cambridge, UK) with a custom-written script. As the rats crossed a central photo-detector, visual cues (VCs) were displayed in pseudo-random sequence on video monitors positioned behind, and parallel to the two reward arms. This is the “VC”, or “central arm PD” event. The VCs were either lit or dim uniform fields. The rat then selected the left or right arm and crossed the Reward arm (Rew) photodetector (PD), triggering reward release from an audible solenoid valve. Crossing the photodetector in the middle of the return arm of the T-maze (Return arm PD; Ret) triggered the visual cue to be turned off. Photodetectors detected task events and triggered cues and rewards via the CED Spike2 script.

### 2.3 Viral vector preparation, injection and immunohistochemistry

The Canine Adenoviral vector (CAV2-PRS-ChR2-mCherry) was produced at the University of Bristol. Details about it and injections procedures appear in Xiang, et al. (2019). Also see Xiang, et al. (2019) for histological and immunohistochemical procedures and neuron characterization.

### 2.4 Electrode and optrode implants

Following VC task pre-training, at least one day before surgery, rats were returned to ad libitum water and food. Surgical procedures and electrode construction are the same as in Xiang, et al. (2019). Moveable tungsten microelectrodes (insulated with epoxylite®, impedance = 2-4 MΩ, FHC Inc, USA) were used for LC recordings. A single microelectrode, or two or three such electrodes glued together was implanted at AP -3.8-4 mm relative to lambda, and ML 1.1-1.2 mm, with a 15° rostral tilt. A stainless steel wire (Teflon coated, diameter=178 µm, A-M systems Inc) implanted in the midbrain area about 1-2 mm anterior to the LC electrode tip served as a fixed LC reference electrode, permitting differential recording. The rat with the virus injection (R328) was implanted with an optrode composed of a tungsten microelectrode (insulated with epoxylite, impedance = 2-4 MΩ, FHC Inc, USA) glued to a 200 µm optic fiber implant with a ferrule (0.37 numerical aperture, hard polymer clad, silica core, multimode, Thorlabs), with tip distances 1 mm apart (the electrode was deeper). The optic fiber implant and optic fiber cables were constructed at the NeuroFabLab (CPN, Ste. Anne Hospital, Paris). Two screws (diameter = 1 mm, Phymep, Paris) with wire leads were placed in the skull above the cerebellum to serve as ground. LC electrodes were progressively lowered under electrophysiological control until characteristic LC spikes were identified (located ∼ 5-6 mm below the cerebellar surface, see Xiang, et al., 2019 for details). For the virus-injected rat, LC spikes could also be identified by responses to laser stimulations (described below). Following implantation, the microelectrode was fixed to a micro-drive allowing for adjustments along the dorsal-ventral axis. The headstage was fixed to the skull with dental cement, and surrounded by wire mesh stabilized with dental cement for protection and shielding. After the surgery, animals were returned to their home cages for at least one-week recovery with ad libitum water and food and regular observation.

### 2.5 Electrophysiological recordings

Rats were then returned to dietary restriction. The movable electrodes were gradually advanced until a well-discriminated LC unit was encountered and then all channels were recorded simultaneously while the rat performed in the T-maze. If no cells could be discriminated, the electrodes were advanced and there was at least a 2 h delay before the next recording session.

For daily online monitoring of LC spikes, pre-amplified signals were filtered between 300-3000 Hz for verification on the computer screen (Lynx-8, Neuralynx, Bozeman, MT, USA) and also transmitted to an audio monitor (audio analyzer, FHC). For recordings, brain signals were pre-amplified at unity gain (Preamp32, Noted Bt, Pecs, Hungary) and then led through a flexible cable to amplifiers (x500, Lynx-8, Neuralynx) and filters (0.1-9 kHz, Lynx-8, Neuralynx). Brain signals were digitized at ∼20 kHz using CED Power1401 converter and Spike2 data acquisition software. The LC unit activity was identified as in Xiang, et al. (2019) by: 1) spike waveform durations ≥0.6 ms; 2) low average firing rate (1-2 Hz) during quiet immobility; 3) brief responses to unexpected acoustic stimuli followed by prolonged (around 1 s) inhibition; 4) for the virus-injected rat (R328), LC units were verified by responses to laser stimulation. A laser driver (Laserglow Technologies, Canada, wavelength 473 nm) was controlled by signals from a stimulator (Grass Technologies, USA, Model SD9). Light intensity from the tip of optic fiber was measured by a power meter (Thorlabs, Germany, Model PM100D). If unit firing was entrained to the pulses with an increased rate (to at least twice the baseline firing rate) averaged over all the stimulations, they were considered to be noradrenergic LC units.

A light emitting diode (LED) was mounted on the cable that was plugged into the headstage. This was detected by a video camera mounted above the T-maze and transmitted to the data acquisition system at a sampling rate of ∼30 Hz for the purpose of position tracking.

### 2.6 Signal processing, spike sorting and data analyses

For off-line spike detection of LC activity in three of the rats, the wide-band signals were converted and digitally high-pass filtered (nonlinear median-based filter). Waveforms with amplitudes passing a threshold were extracted, and then subjected to principal component analysis (PCA). All of these processes were performed with NDManager (Hazan, et al., 2006). Spikes were sorted with a semi-automatic cluster cutting procedure combining KlustaKwik (KD Harris, http://klustakwik.sourceforge.net) and Klusters (Hazan, et al., 2006). Spikes with durations less than 0.6 ms were rejected. In one rat (R311), the LC signal was filtered from 300-3000 Hz during recording, and the spike sorting was performed with Spike2 software (which employs a waveform template-matching algorithm). Most data analyses were performed using Matlab (R2010a) with the statistical toolbox FMAToolbox (developed by M. Zugaro, http://fmatoolbox.sourceforge.net) and scripts developed in the laboratory as well as some statistical analyses performed with Microsoft© Excel©. Phase was computed with the ‘Phase’ function of the FMAToolbox, employing the Hilbert transform. To characterize periods with infra-slow rhythms, a criterion for salient phase-locking to task events was established as when the SEM range of LFP phase was less than 0.75*π radians (cf., Figure 4, middle column). This is termed “regular phase-locking”. Sessions tallied for phase-locking of LC neurons to infra-slow rhythms were included only if they had at least 1000 LC neuron spikes.

## 3 Results

### 3.1 LC neuron phase-locking to prefrontal and hippocampal infra-slow rhythms

Infra-slow rhythms were readily apparent in visual inspections of hippocampal (Hip) and prefrontal cortical (Pfc) local field potentials (LFPs) (Fig. 1A). These were rendered more salient by filtering the signal in a 0.1-1.0 Hz window (Fig. 1B). We applied an amplitude threshold to examine data from those periods when the infra-slow rhythm amplitude was elevated (Fig. 1C). We applied an amplitude threshold to examine data from those periods when the infra-slow rhythm amplitude was elevated (Fig. 1C). This was intended to limit analyses to those periods when the infra-slow oscillation was sufficiently robust, thus avoiding possibly spurious computations of phase angle from low amplitude oscillations. In each of five rats, LC neurons were phase-locked to Pfc (e.g., Figs. 1D and 2), as well as Hip infra-slow LFP rhythms. The incidence of phase-locking of the LC neurons in sessions with regular phase-locking of the infra-slow rhythms to task events was 20 out 23 for Hip LFP and 15/20 for Pfc LFP (Rayleigh test, p<0.05; for histology, see Fig. 2 of Xiang, et al., 2019). In the animal where noradrenergic LC neurons were identified optogenetically (see Methods), all were phase-locked to the infra-slow rhythms (n=8 for both Pfc and Hip; Rayleigh test, p<0.05).

**Figure 1.**
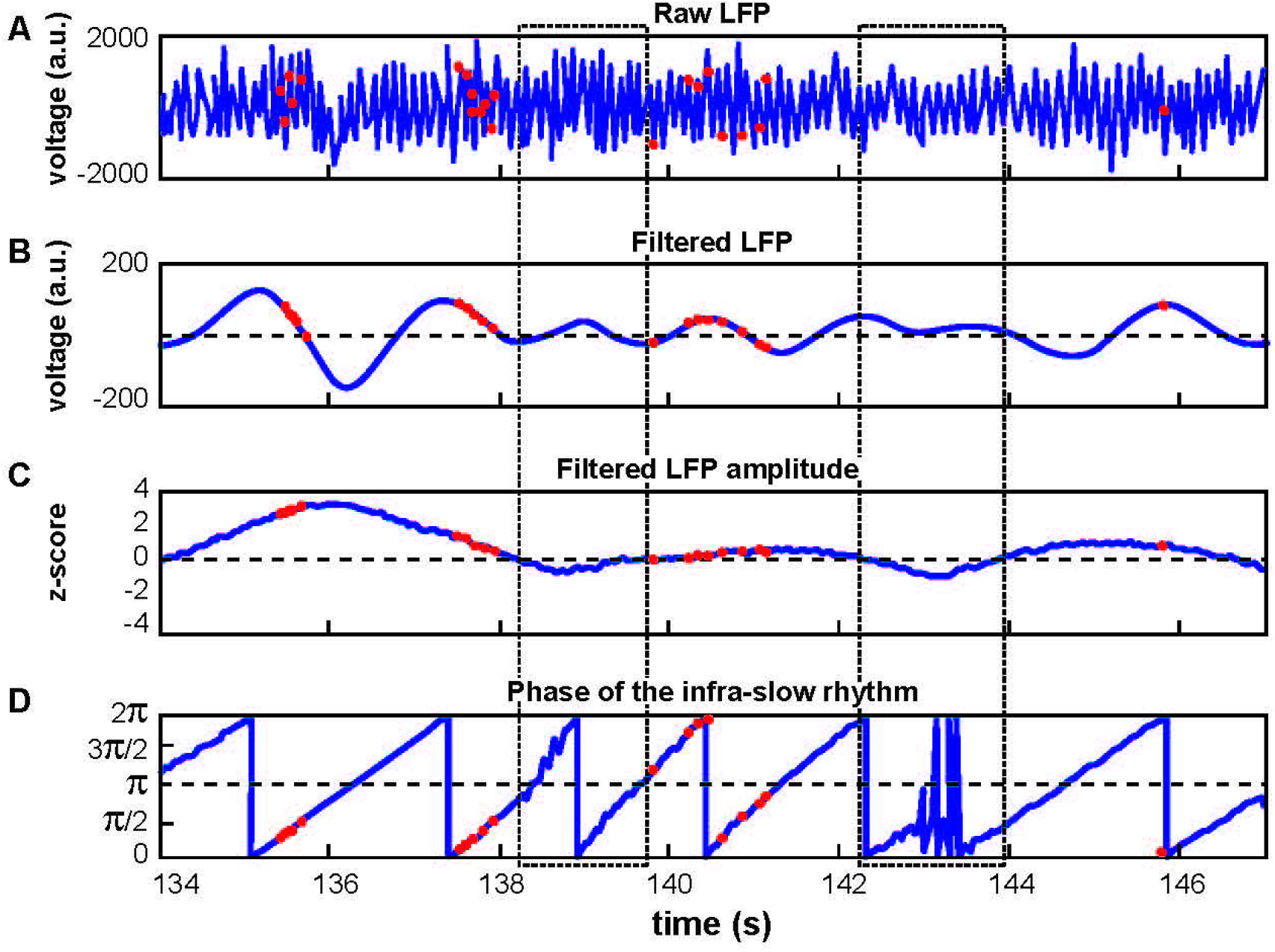
Calculation of LC spike phase relative to Hip or Pfc LFP. A) Unfiltered signal of a hippocampal recording, with theta oscillations dominating. B) The signal from A band-pass filtered at 0.1-1.0 Hz. Red dots indicate LC neuron action potentials in all panels. C) The amplitude of the signal in B was z-scored. Low amplitude oscillations were excluded from analyses according to a selected criterion of z≤0 (excluded zones are demarcated by the dotted rectangles). D) Phase of the filtered signal in B. Note that the LC spikes generally occur at phases between 0 and π/2 radians in this example. The discontinuities near 138.5 and 143 s correspond to excluded data, where phase could not be computed reliably.

**Figure 2.**
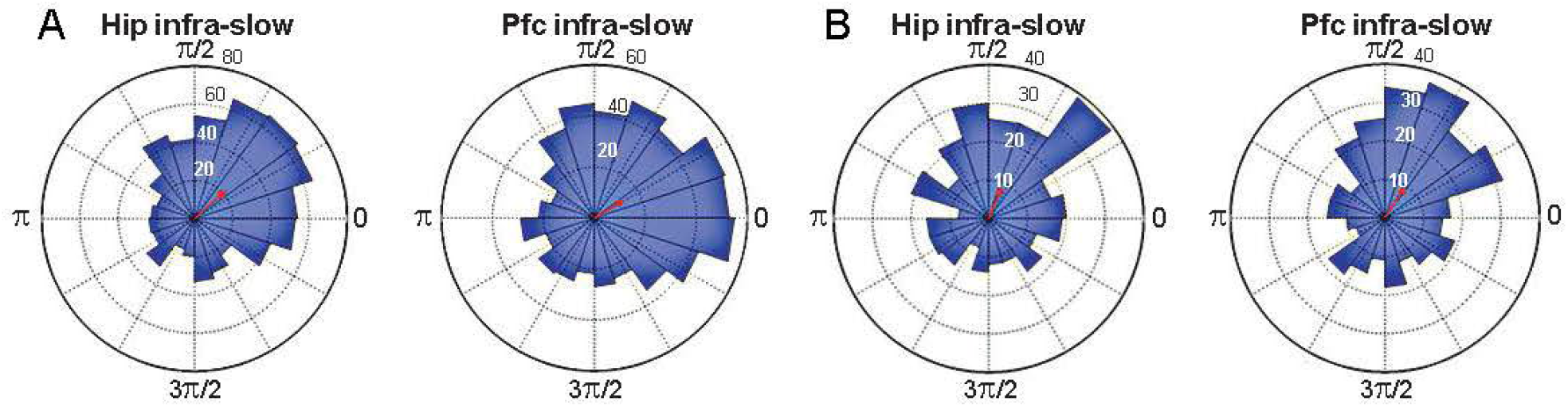
Spike phase-locking to infra-slow rhythms from two example LC neurons (A and B). Radius values are spike counts. Red arrows represent resultant vectors.

The modal preferred infra-slow phase among these neurons was 0.35*π radians for Hip infra-slow and 0.15*π radians for Pfc infra-slow (p<0.05, Rayleigh test; not shown).

### 3.2 Prefrontal and hippocampal infra-slow rhythms are synchronized to maze events

The infra-slow rhythms were phase-locked to task-relevant positions on the maze (Fig. 3B; Supp. Fig. 1B). To quantify this phase-locking, the mean (± SEM) phase of the rhythm was plotted in peri-event time color plots (see Fig. 4 and Supp. Fig. 1A for examples) over all trials in 57 sessions from eight rats (including the five with LC recordings). Infra-slow rhythms were phase-locked to the reward arm photodetector crossing (Rwd) in 51 of the recording sessions for Hip, and 46 sessions for Pfc (see Table 1). The other maze events had fewer incidences of regular phase-locking (Pfc return arm photodetector crossing, or Rtn: 11; Pfc central arm visual cue onset PD, or VC: 18; Hip Rtn: 18; Hip VC: 20). The mean phases at the respective PD crossings (in those cases when SEM≤0.75*π radians there) were 0.70*π and 0.24*π radians for Pfc and Hip Rtn, 0.25*π and 0.22*π radians for Pfc and Hip Rwd, and 0.19*π and 0.01*π radians for Pfc and Hip VC. The root-mean-square differences between Pfc and Hip mean phase (calculated pairwise by session) at the respective PD crossings were 0.14*π, 0.13*π and 0.12*π rad. The regular phase-locking could last from less than one to over 2.5 successive rhythmic cycles (Supp. Figs. 1, and 2, Table 1) and could continue from one event to the next (Fig. 4, Supp. Fig. 1). For PL Rwd and Hip Rwd, 30 and 37 sessions had durations of regular phase-locking lasting one or more cycles, respectively. These permitted quantification of the temporal duration of the cycles, which ranged from 2.0 to 2.6 s, the equivalent of 0.4 to 0.5 Hz. In the six cases of Rwd PD phase-locking which had a second complete cycle, the mean of the first was 2.3 s, while the second was lower, 2.0 s (pairwise t-test, p=0.0009, df=5). Thus, these are not regular periodic oscillations, but, rather, this is consistent with phase-locking to task events. Pfc and Hip infra-slow rhythms sometimes resembled one another (e.g., Fig. 2). To compare them, sessions were classified as having Pfc and Hip regular phase-locking in the following ranges of cycles (see Table 1). In 17 of the 57 sessions, these numbers of cycles were different between Pfc and Hip for VC, Rwd and/or Rtn (e.g., Supp. Fig. 2). This indicates that it is unlikely that Pfc and Hip infra-slow rhythms are related by volume conduction, and suggests that they could be independently generated.

**Figure 3.**
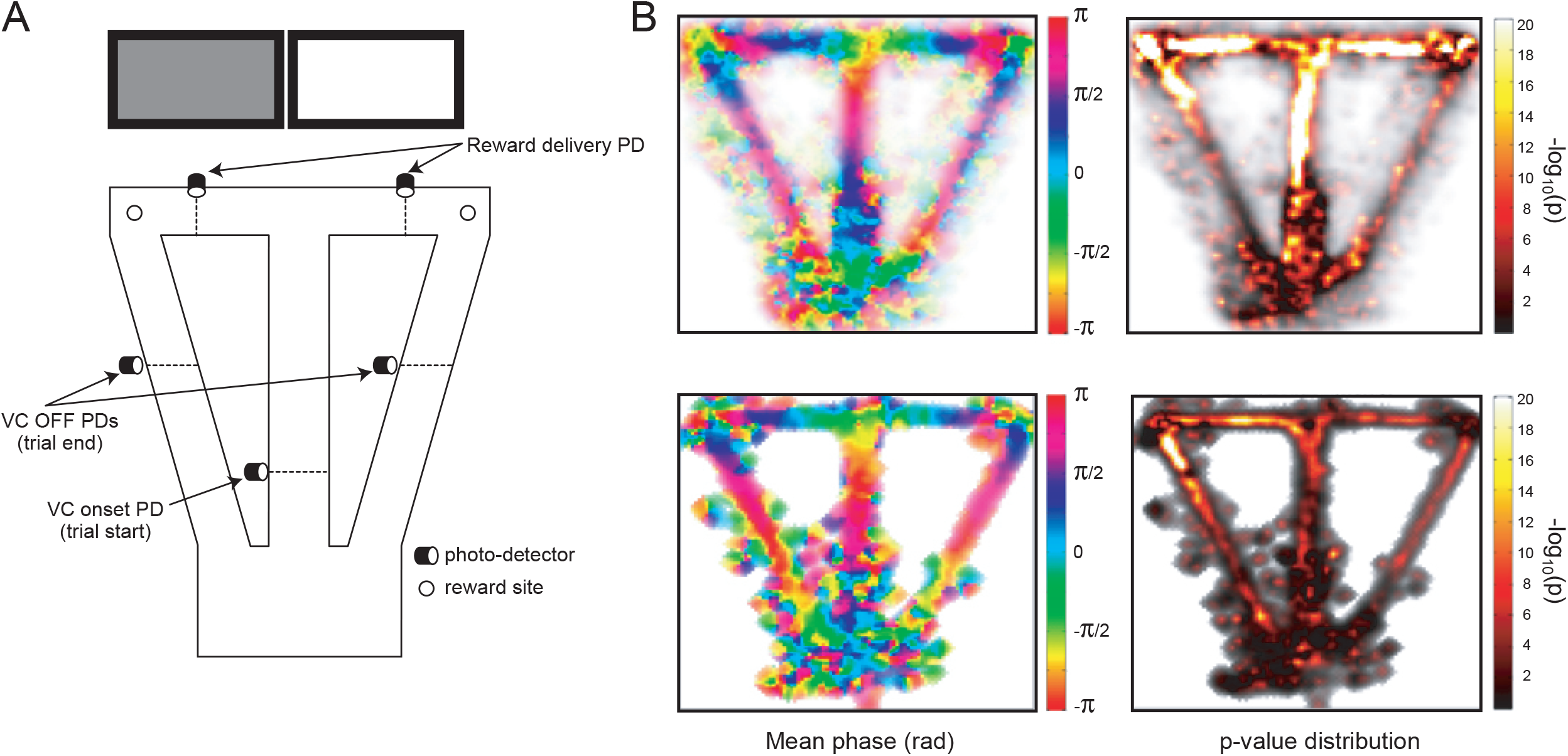
A) The automated behavioral task. When the trained rat crosses the central arm photodetector (VC onset PD), this triggers one of the two cue screens behind the reward arms to be lit in pseudo-random sequence. Crossing the appropriate reward delivery PD triggers a drop of sweetened water to arrive at the corresponding reward site. Crossing the VC OFF PD’s on the return arms triggers the lit screen to be turned off. These three photodetector events are used to synchronize activity in other Figures. B) Distribution of mean phase (left) and p-values of phase-locking (right; Rayleigh test) for Pfc infra-slow oscillations in pooled data from multiple sessions (top), and in an example session (bottom).

**Figure 4.**
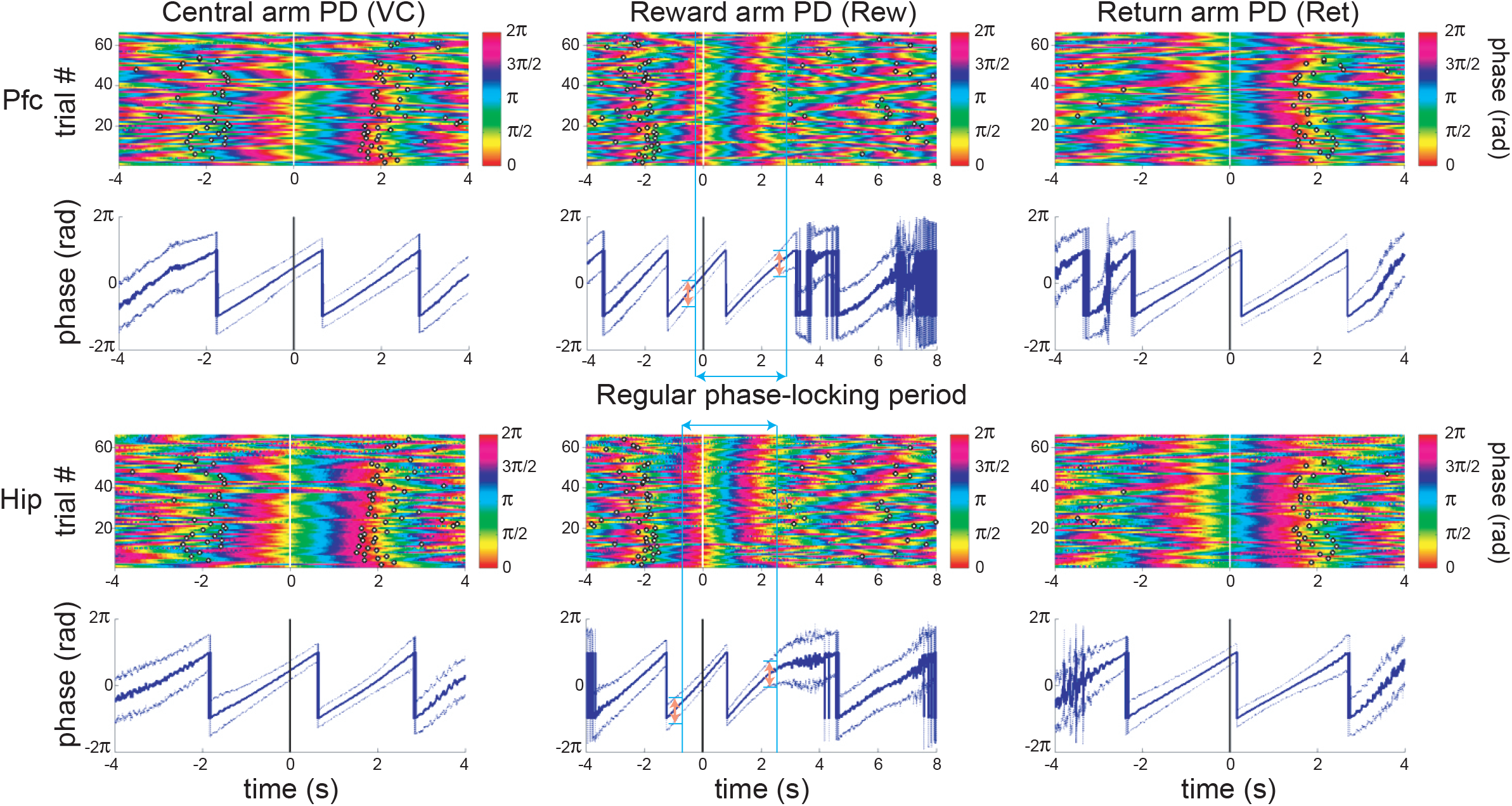
An example of simultaneous recordings of Pfc and Hip LFP infra-slow oscillations phase-locked to principal maze events, the PD crossings (at time zero). Each row of the color plots corresponds to a single trial. The phase of the infra-slow LFP is color-coded. Black rings correspond to the PD crossing prior to (left) or after (right) the event at zero for each plot. Note that the time scales vary among the events, in order to display prior and subsequent PD’s. The dark traces in the 2^nd^ and 4^th^ rows show mean phase while the lighter shaded upper and lower lines are ±SEM. In the middle column, the blue vertical bars and blue double-headed arrow illustrate the calculation of the range of regular phase-locking (defined here as the period with the criterion of SEM range<0.75*π radians; pink double-headed arrows). Here, desynchronization (zones with wider SEM ranges) and discontinuities in the mean phase result from inter-trial variability in speed and distance from the synchronization point. (PD - photodetector crossing). This is from the same session as the recording in Fig. 3B.

**Table 1.**
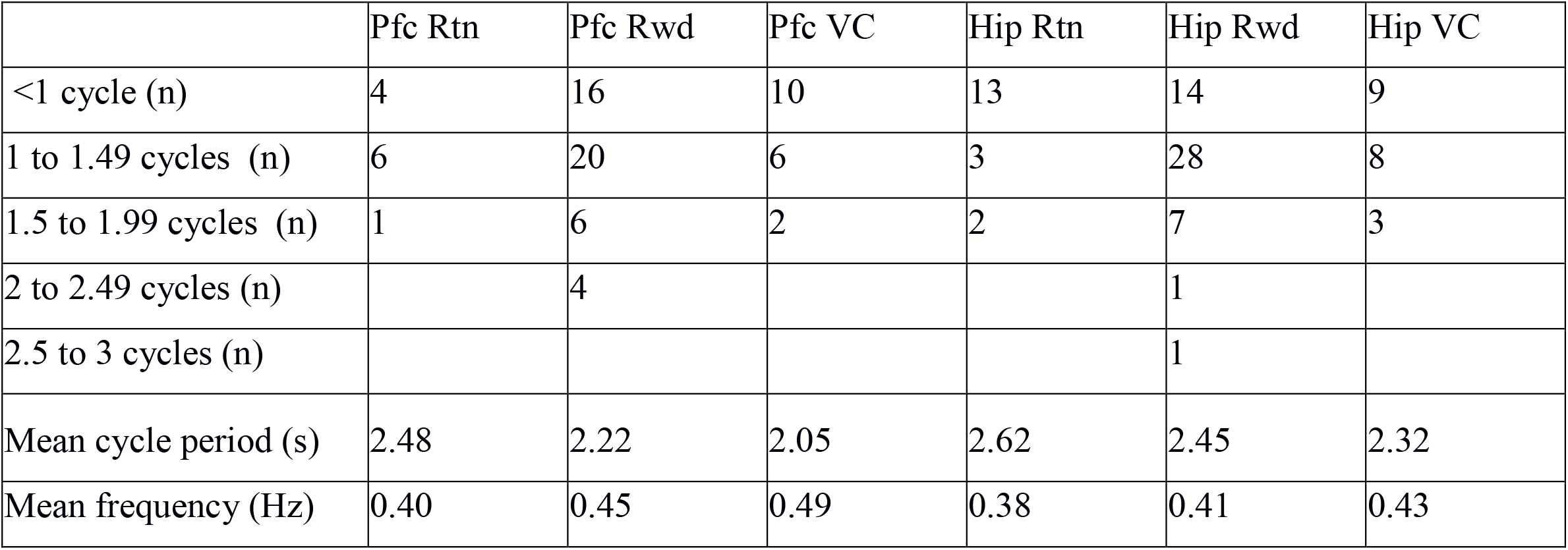
Characterization of periods in sessions with regular phase-locking (SEM≤0.75*π radians) of infra-slow LFP oscillations in prefrontal cortex (Pfc) and hippocampus (Hip) to task events. In the six cases of two or more cycles, only data from the first cycle were counted for mean cycle period and frequency. Cycles are only counted in the period from the previous trial event to the next one, even though successive cycles could extend before or after (cf., Fig. 4, Supp. Fig. 1).

|  | Pfc Rtn | Pfc Rwd | Pfc VC | Hip Rtn | Hip Rwd | Hip VC |
| --- | --- | --- | --- | --- | --- | --- |
| <1 cycle (n) | 4 | 16 | 10 | 13 | 14 | 9 |
| 1 to 1.49 cycles (n) | 6 | 20 | 6 | 3 | 28 | 8 |
| 1.5 to 1.99 cycles (n) | 1 | 6 | 2 | 2 | 7 | 3 |
| 2 to 2.49 cycles (n) |  | 4 |  |  | 1 |  |
| 2.5 to 3 cycles (n) |  |  |  |  | 1 |  |
| Mean cycle period (s) | 2.48 | 2.22 | 2.05 | 2.62 | 2.45 | 2.32 |
| Mean frequency (Hz) | 0.40 | 0.45 | 0.49 | 0.38 | 0.41 | 0.43 |

The infra-slow rhythms were regularly phase-locked to two (in 24 sessions), or even all three (in 8 sessions) different task events. Thus, they were not linked to any specific task-related behavior. To test whether infra-slow rhythms were triggered by rapid head movements, regression analysis compared the onset of regular phase-locking and times of peak acceleration, or deceleration around the Rwd PD crossings, and were not significant (r^2^=0.034, p=0.49 and r^2^=0.0056, p=0.80 respectively; df=15; see Supp. Fig. 3). In Xiang et al. (2019), we showed that LC neurons fire more during accelerations. Indeed, the periods with the greatest increase in LC activity were not those most frequent for the start of regular phase-locking (i.e., reset) of the infra-slow rhythm; rather phase-locking occurred most frequently to Rwd PD crossing (see above), where no consistent accelerations occurred (see Supp. Figs. 3 and 4). These results indicate it is unlikely that Pfc and Hip infra-slow rhythms are due to a biomechanical artifact, for example, from locomotion or head rocking.

### 3.3 Coordination of neuronal activity across time scales

In the four sessions where Hip and Pfc neurons could be discriminated from the LFP electrodes, most were also modulated by infra-slow rhythms (Pfc LFP modulated 6/12 Pfc units and 8/11 Hip units; Hip LFP modulated 8/12 Pfc units and 8/11 Hip units; Rayleigh test p<0.05). The LC neurons could have relatively consistent phases with respect to the two infra-slow rhythms (not shown). LC neurons could be phase-locked to oscillations in the delta frequency range (1-4 Hz) in Pfc (n=15/37) and Hip (11/37) as well as theta (5-10 Hz; 7/37 and 5/37 respectively) for Pfc and Hip (see Supp. Fig. 5). While phase-locking of LC neurons to gamma (40-80 Hz) was rare (n=2 for both structures’ LFPs), the infra-slow rhythm did modulate the amplitude of gamma oscillations at 35-45 Hz in hippocampus and prefrontal cortex (Fig. 5).

**Figure 5.**
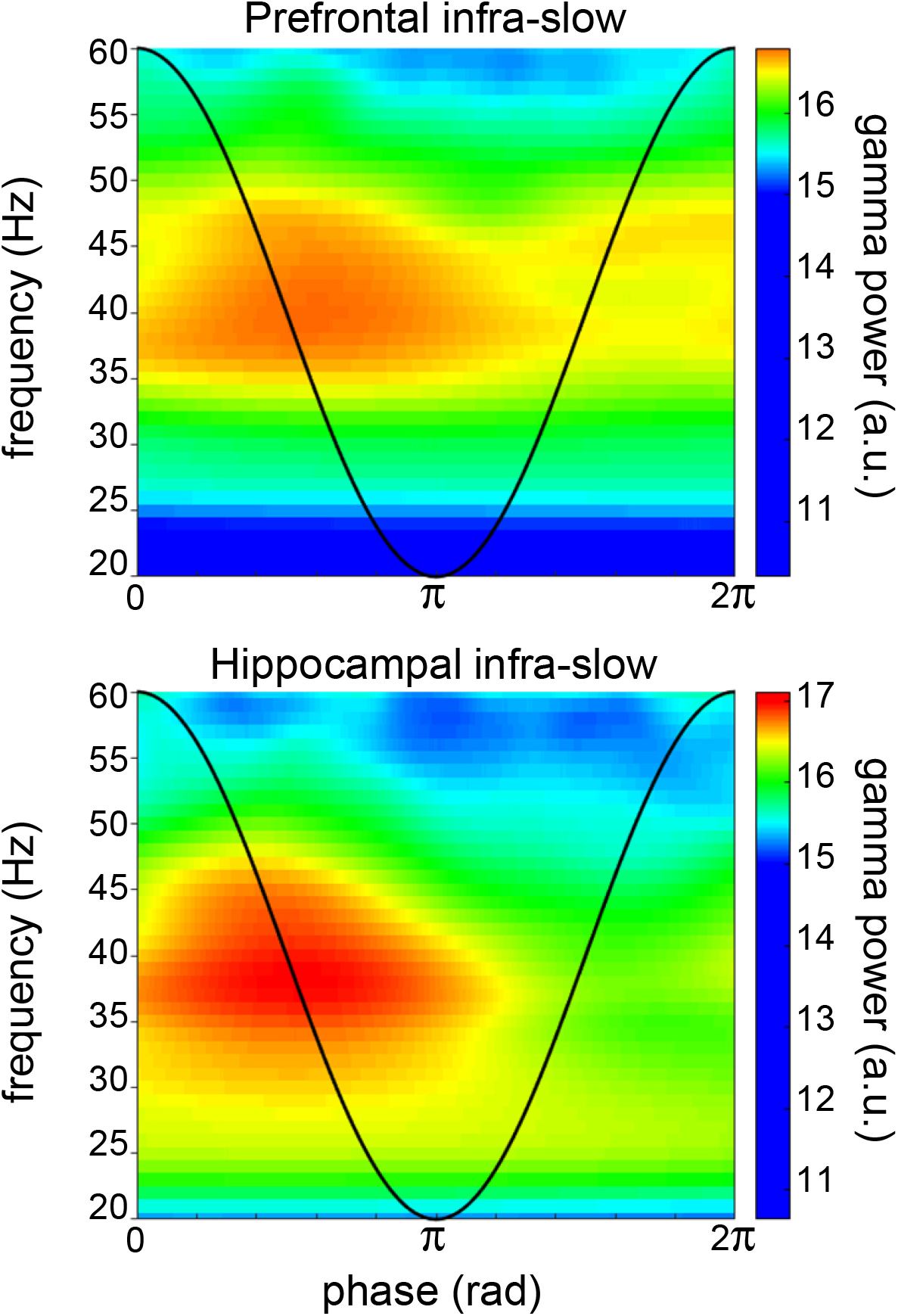
Example of infra-slow modulation of gamma rhythm LFP in Pfc (top) and Hip (bottom). The black trace schematically illustrates the phase of the infra-slow rhythms. Gamma power was elevated between the phases of zero and pi radians of the infra-slow rhythm, while this was not apparent at other frequencies. a.u. – arbitrary units.

## 4 Discussion

LFP oscillation cycles on the order of 0.4 Hz in prefrontal cortex and hippocampus were phase-locked to task events at crucial points on the maze. Successive cycles had different cycle lengths, indicating that, if they are indeed periodic oscillations, their phase can reset to salient events. Simultaneous recordings in prefrontal cortex and hippocampus could have different cycle lengths as well, while still phase-locking to task events. This would seem to exclude any single structure from entraining these independent rhythms simultaneously. This also would rule out a contribution of volume conduction. The intriguing issue of the origin of these rhythms merits further investigation. Most of the LC neurons were phase-locked to these infra-slow prefrontal cortical and hippocampal LFPs, including all of the optogenetically identified noradrenergic neurons. Hippocampal and prefrontal units were also phase-locked to the infra-slow oscillations. While the number of LC neurons recorded may appear low, this is typical for the rare chronic recording studies of this structure in behaving rats, likely because its diminutive dimensions and deep location render accurate electrode placements challenging.

This is consistent with previous work showing neuronal activity adapting to the time scale of behavioral events. For example, in behavioral tasks with delays, several brain structures show “time cell” activity: neurons with sequential “tiling” activity lasting on the order of several seconds. These periods can expand or contract depending upon the duration of task-imposed intervals (MacDonald, et al., 2011). We speculate that this infra-slow rhythm may originate in the hippocampal-prefrontal system since neuro-physiological activity there tracks time intervals on the order of several seconds based upon regularities in temporal structure of behavioral or environmental events.

Steriade, et al. (1993) observed infra-slow (0.3-1.0 Hz) rhythms in neocortical activity in anesthetized and naturally sleeping cats. Eschenko et al (2012) showed that LC neuronal activity in sleeping rats is synchronized with the sleep infra-slow wave cycle (1 Hz) and is out of phase with Pfc neuronal activity. Similarly, in rats under ketamine anesthesia, there is a negative correlation between activity of LC NE neurons and prefrontal neurons, when neuron activation oscillates at ∼1 Hz (Sara and Hervé-Minvielle 1995; Lestienne, et al. 1997). Furthermore, when the latter authors pooled their LC recording data, they were significantly phase-locked to cortical LFP delta oscillations. While these infra-slow cycles of UP-DOWN state transitions are not generally observed in awake animals, this does demonstrate that LC can fire rhythmically, and that these structures can coordinate their activity at this time scale. Furthermore, in rats under urethane anesthesia, Totah, et al. (2018) found that the firing rate of locus coeruleus neurons oscillates at 0.4-0.5 Hz. And, in head-fixed awake mice, cortical noradrenergic axons exhibited rhythmic Ca^2+^ activity at 0.5–0.6 Hz (Oe, et al., 2020). Thus, the LC could also be associated with the Pfc-Hip in the origin, maintenance and communication of behaviorally relevant infra-slow rhythms in the brain. Further work is required to elucidate the respective roles of these structures in these processes.

In the awake state, there is evidence for infra-slow neural processing although this was not observed as rhythms per se. Molter, et al. (2012) observed a 0.7 Hz modulation of the power of theta rhythm recorded in rat Hip. This 0.7 Hz modulated Hip neuronal activity during sleep, as well as during behavior in a maze, a running wheel, and an open field. Positions on a figure-8 maze corresponded to specific phases of this modulatory rhythm, similar to the infra-slow rhythm recorded here. (Their filter settings excluded 0.7 Hz rhythms and thus this could not be directly measured in that work.) In Molter et al. (2012), the 0.7 Hz modulation of the power of the theta infra-slow modulation was locked at π radians to junction points in the maze (their Figure 7B), where accelerations might be expected. However, they found no overall correlation between phase and acceleration. Halgren et al. (2018) observed a rhythm at less than 3 Hz generated in the superficial layers of the cerebral cortex in awake humans. The phase of this rhythm reset to infrequent tones in their oddball task, similar to the reset of the infra-slow rhythm here in relation to salient task events.

Villette, et al. (2015) used calcium imaging to observe CA1 pyramidal cells in head fixed mice moving in the dark on a non-motorized treadmill. They found that different neurons fired sequentially in cycles at the same time scale as the infra-slow oscillations observed here. Furthermore, the cycles could occur singly, or consecutively in groups of two or three. The authors interpreted this as representing an intrinsic metric for representing distance walked. This resembles time cell activity (Pastalkova et al., 2008; MacDonald et al., 2011) evoked above, where the length of the cycle extends to the time scale of the ongoing task (Kraus, et al., 2013; Ravassard, et al., 2013). The 2 to 5 s durations of the cycles in the Villette, et al. (2015) study may represent a default value since their task had no temporal structure. This is on the order of the time scale of the infra-slow rhythm recorded here, and the variable numbers of cycles they observed might flexibly adapt to the positions of task-relevant events to lead to the results found here.

The present observations of phase-locking of LC neurons to infra-slow rhythms in hippocampus could ostensibly be due to independent synchrony of the infra-slow rhythms and the LC neurons to task events. However, the LC neurons showed phase preferences in the infra-slow rhythms in data pooled over multiple task events. We did not observe any simple relation between infra-slow rhythms and motor events (e.g., as we showed for LC neurons with acceleration or deceleration by Xiang, et al., 2019), since regular phase-locking could start before (Supp. Fig. 1) or after the same task events in different sessions (not shown), and continue over periods including a variety of behaviors.

The phase-locking of LC neurons to infra-slow rhythms in Hip and Pfc, as well as to oscillations in the delta, theta and gamma frequency bands could reveal coordinated neuronal processing within a unified temporal framework (cf., Totah, et al., 2018a). The scale of this corresponded to the temporal and spatial regularities characterizing the current behavioral patterns. Cross-frequency coupling could serve as a mechanism to link processing at different time scales. This could facilitate both ‘Communication through coherence’ (CTC, Bosman et al., 2012; Fries, 2005) and ‘Binding by synchrony’ (Eckhorn, et al., 1990; Engel, et al., 1999; Buehlmann and Deco, 2010). Thus, infra-slow rhythms would serve as a scaffold to link the time scales of dynamics of neuronal processes to those of behavior and cognitive processes. Noradrenaline, released by LC neurons in concert with the infra-slow rhythm, would participate in synchronizing or resetting those brain networks underlying behavioral adaptation to these events (Bouret and Sara, 2005; Sara and Bouret, 2012).

## 5 Competing interests

The authors declare that the research was conducted in the absence of any commercial or financial relationships that could be construed as a potential conflict of interest.

## 6 Author contributions

S.I.W. and L.X. designed the experiments; S.J.S. and L.X. developed and implemented the LC optogenetics and recordings; L.X. and H.Y.G. performed the experiments; L.X., S.I.W., R.T., A.H. and S.J.S. designed the analyses; L.X., R.T., and A.H. performed the analyses; S.J.S., S.I.W., and L.X. wrote the paper. All authors approved of the final version of the manuscript.

## 7 Funding

L.X. was supported by a fellowship from the China Scholarship Council (CSC). The Labex Memolife and Fondation Bettencourt Schueller provided support.

## 8 Acknowledgements

Thanks to Professor Anthony E. Pickering for providing the virus and related advice. Thanks to Dr. Michaël Zugaro for helpful suggestions and help with analyses and computing, Drs. A Sirota and X Leinkugel for helpful discussions, and France Maloumian for help with figures.

## 9 Contribution to the field statement

Periodic oscillations of excitability coordinate neuronal activity within and between brain structures for perception, cognition and goal-directed behavior, processes implicating noradrenergic activity in forebrain circuits. To better understand the link between the time scales of behavior (on the order of seconds) and underlying neuronal processing (on the order of milliseconds), we recorded phase-locking of neurons in the noradrenergic locus coeruleus to brain oscillations in rats performing in a maze. Most neurons synchronized with hippocampal and prefrontal cortical infra-slow (∼0.4 Hz) rhythms. The infra-slow rhythms phase-locked to principal events in the maze, and thus were not strictly periodic. They modulated the amplitude of gamma rhythms, known to coordinate neuron activity, and thus could provide a scaffold linking behavior to neuronal activity.

## Notes

### Competing Interest Statement

The authors have declared no competing interest.

### Summary of Updates

Revision contains corrections and clarifications and a new Supp. Fig 5. Repetitions of methods have been shortened or removed, referring to Xiang et al (Sci. Rep., 2019).

