## Supplementary material for "Locus cœruleus noradrenergic neurons phase-lock to prefrontal and hippocampal infra-slow rhythms that synchronize to behavioral events": Supp. Fig. 1

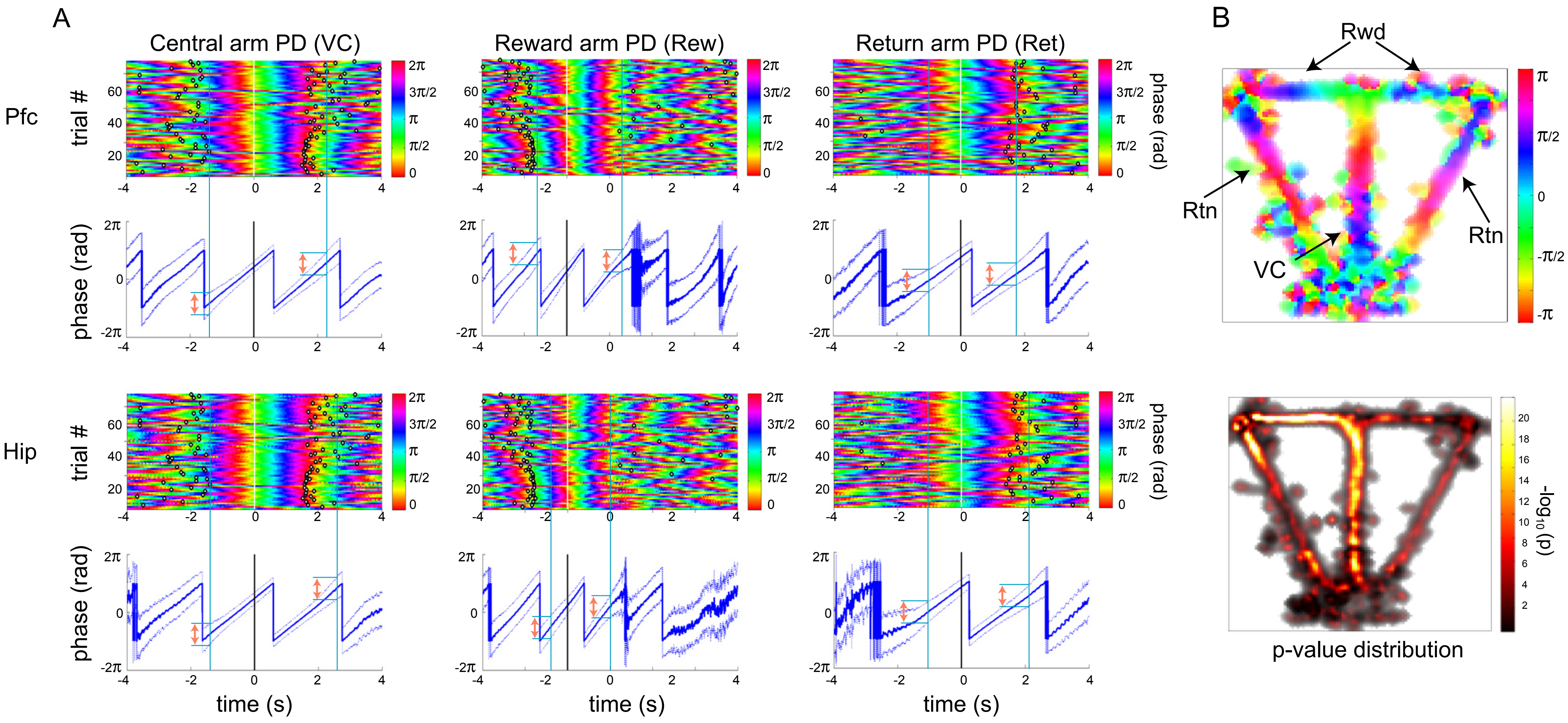


Supplementary Figure 1. The infra-slow cycles could continue through sequential phases from one task event to the following one. Plots are in same format as in Figure 4. A) Note that regular phase-locking in the left column (VC) starts prior to zero just after the Rtn PD (black dots prior to zero) and continues up to Rwd PD (black dots after zero). The other columns are comparable. B) Mean infra-slow phase distribution for this session (as in Fig. 3B) also shows the continuity of the phase distribution on maze. Arrows show photodetector positions. (Note that colors indicating mean phase are different at the respective photodetector types, and that the color codes in the scales of A and B are not the same.)
