## Supplementary material for "Locus cœruleus noradrenergic neurons phase-lock to prefrontal and hippocampal infra-slow rhythms that synchronize to behavioral events": Supp. Fig. 2

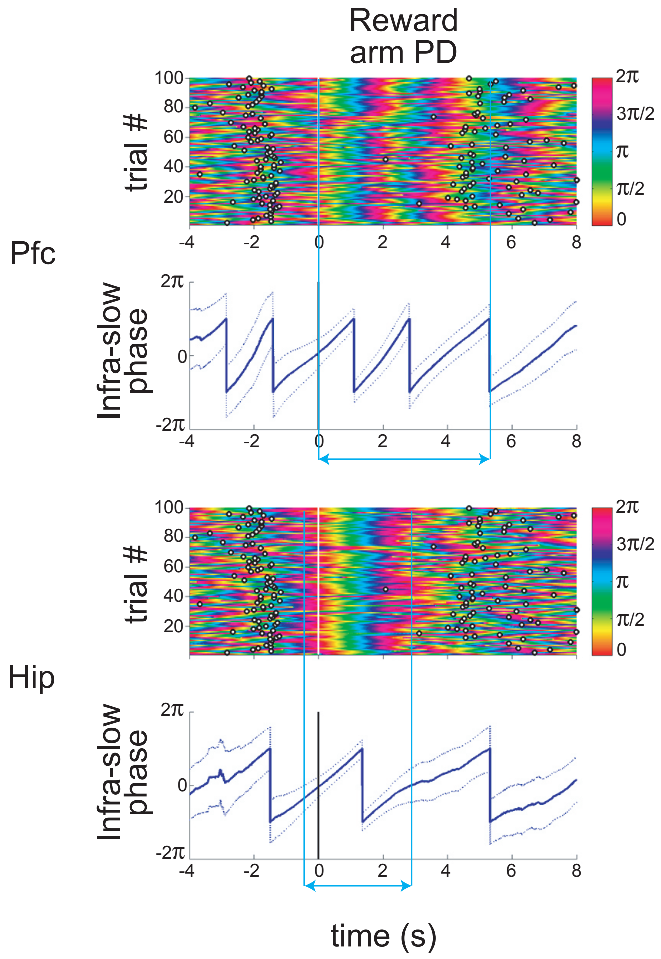


Supplementary Figure 2. Infra-slow rhythms recorded simultaneously in Hip and Pfc can have different cycle lengths. Note that regular phase-locking to Rwd extends for about 2.5 infra-slow cycles in Pfc, but only about 1 cycle for Hip. The durations of the cycles are about 2.1 s (corresponding to 0.48 Hz) for the first Pfc Rwd cycle, 1.9 s for the second (0.53 Hz), and 3.1 s for Hip Rwd (0.32 Hz). (Same format as Figure 4.)
