## Supplementary material for "Locus cœruleus noradrenergic neurons phase-lock to prefrontal and hippocampal infra-slow rhythms that synchronize to behavioral events": Supp. Fig. 3

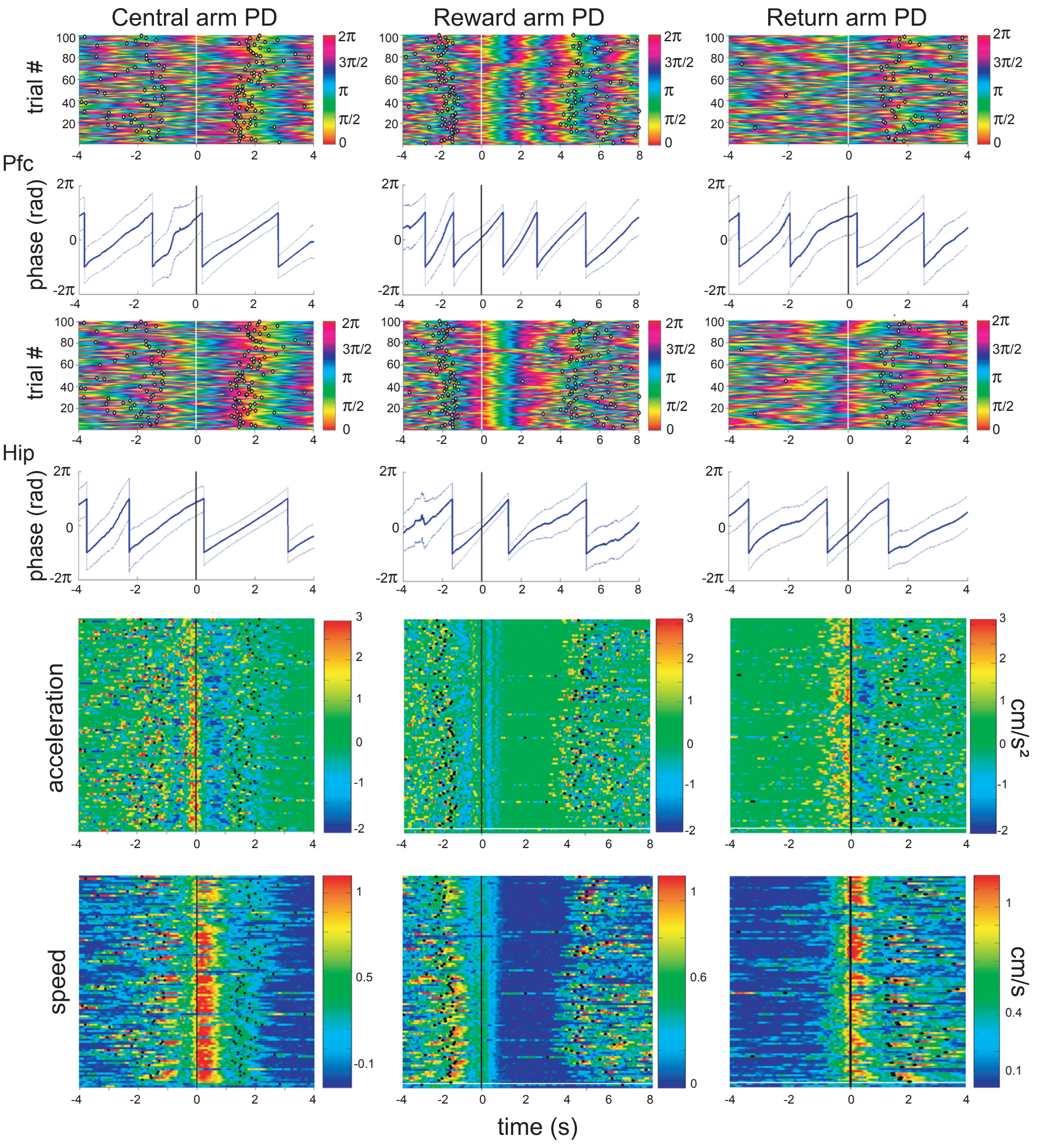


Supplementary Figure 3. Example of lack of a clear relation between speed, acceleration and infra-slow phase. Acceleration increases before central and return arm PD crossings, with speed increasing afterwards. But, the phase is π radians for the former and -0.2*π radians for the latter.
