## Supplementary material for "Locus cœruleus noradrenergic neurons phase-lock to prefrontal and hippocampal infra-slow rhythms that synchronize to behavioral events": Supp. Fig. 4

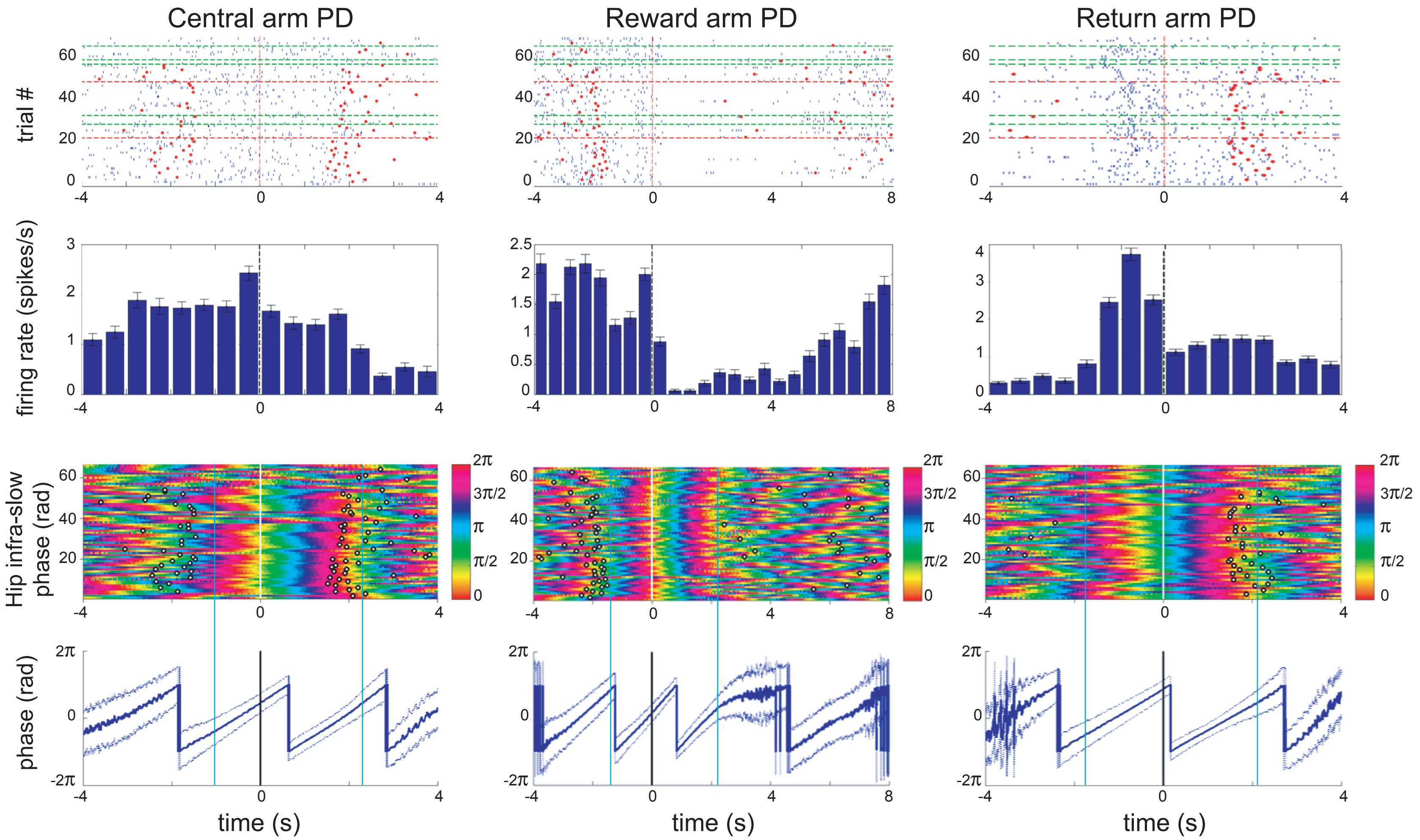


Supplementary Figure 4. Example of infra-slow rhythms persisting during periods of low activity in an LC neuron. Top) Example of LC neuron activity in relation to task events for the cell recorded in the session of Figure 4 (the infra-slow data is reproduced here). This neuron's activity was also shown in the middle row of Figure 4 of Xiang, et al. (2019). Red dots to the left of zero correspond to the previous task event, and those to the right indicate the timing of the subsequent event. Note that the regular phase-locking of infra-slow oscillations at the Reward arm PD continues in the interval (0 s, 2 s), while the LC neuron is inactive. Colored horizontal lines in top row correspond to changes in rules or strategies of the rat (cf., Xiang, et al., 2019), which had no apparent relation to the infra-slow rhythms. Second row) Histogram summing activity in the histograms above. Bottom two rows) Hippocampal infra-slow rhythms for the same session. Vertical blue bars delimit the periods with regular phase-locking.
