## Supplementary material for "Locus cœruleus noradrenergic neurons phase-lock to prefrontal and hippocampal infra-slow rhythms that synchronize to behavioral events": Supp. Fig. 5

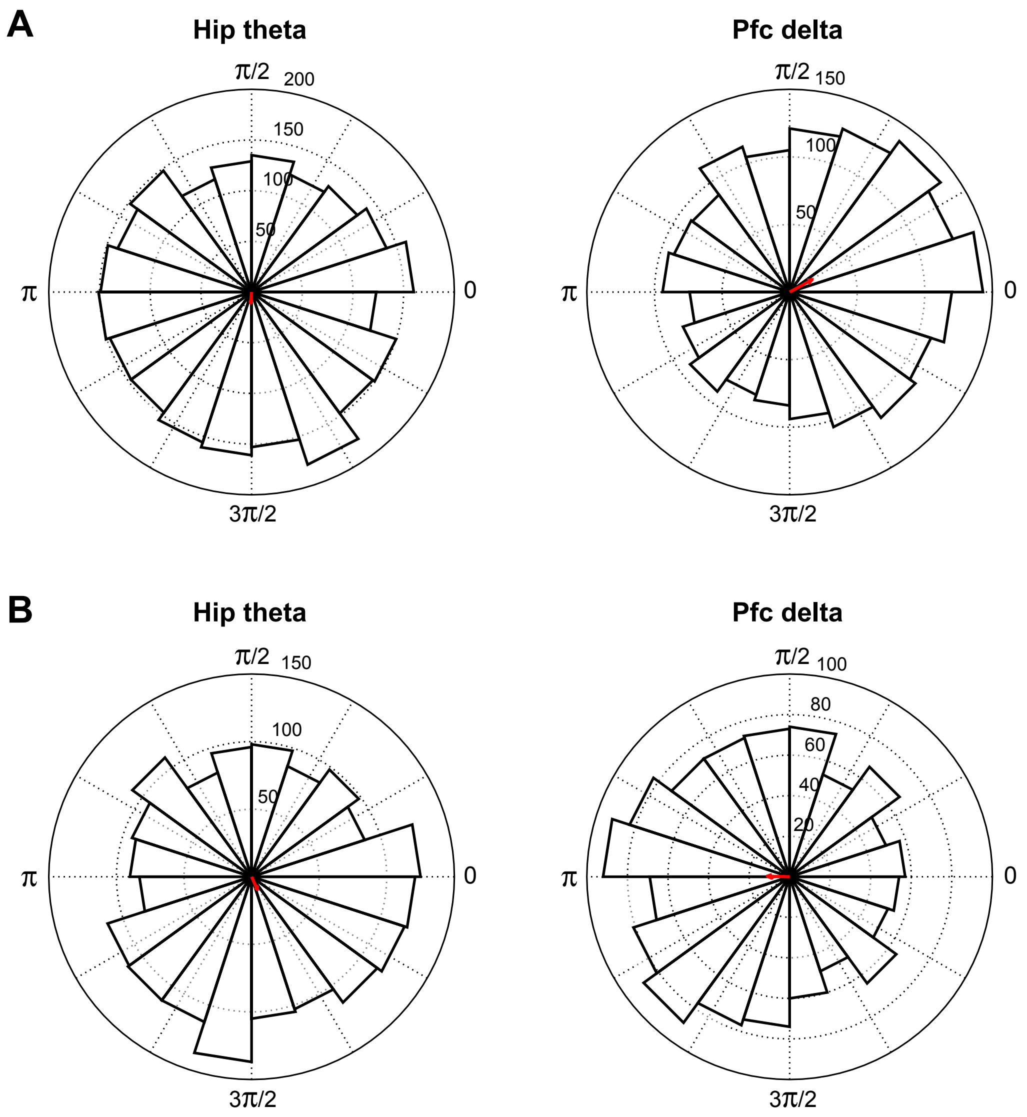


Supplementary Figure 5. Examples of phase-locking of LC neurons to hippocampal theta (filtered 5-10 Hz) and prefrontal delta (filtered 1-4 Hz) rhythms. A and B are two different neurons from two different rats. Same format as Figure 2. A) Hip theta: p=2.6e-4, resultant vector phi=4.6 radians; Pfc delta: p=4.2e-15; phi=0.5 radians, B) Hip theta: p=6.2e-5, resultant vector phi=5.1 radians; Pfc delta: p=7.4e-8; phi=3.1 radians.
